# The unfolded protein response regulates pathogenic development of *Ustilago maydis* by Rok1-dependent inhibition of mating-type signaling

**DOI:** 10.1101/808717

**Authors:** Lara Schmitz, Melina Ayaka Schwier, Kai Heimel

**Author notes:** Address correspondence to Kai Heimel.

## Abstract

Fungal pathogens require the unfolded protein response (UPR) to maintain protein homeostasis of the endoplasmic reticulum (ER) during pathogenic development. In the corn smut fungus *Ustilago maydis*, pathogenic development is controlled by the *a* and *b* mating-type loci. The UPR is specifically activated after plant penetration and required for efficient secretion of effectors and suppression of the plant defense response. The interaction between the UPR regulator Cib1 and the central developmental regulator Clp1 modulates the pathogenic program and triggers fungal colonization of the host plant. By contrast, when activated before plant penetration, the UPR interferes with fungal virulence by reducing expression of *bE* and *bW*, the central regulators of pathogenic development encoded by the *b* mating-type locus. Here we show that this inhibitory effect results from UPR-mediated suppression of the pheromone response pathway upstream of the *b*-regulatory network. UPR activity prompts dephosphorylation of the pheromone-responsive MAPK Kpp2, reducing activity of the pheromone response factor Prf1 that regulates expression of *bE* and *bW*. Deletion of the dual specificity phosphatase *rok1* fully suppressed UPR-dependent inhibition of Kpp2 phosphorylation, formation of infectious filaments and fungal virulence. Rok1 determines activity of mating-type signaling pathways and thus the degree of fungal virulence. We propose that UPR-dependent regulation of Rok1 aligns ER physiology with fungal aggressiveness and effector gene expression during biotrophic growth of *U. maydis* in the host plant.

**Importance:** The unfolded protein response (UPR) is crucial for ER homeostasis and disease development in fungal pathogens. In the plant pathogenic fungus *Ustilago maydis*, the UPR supports fungal proliferation *in planta* and effector secretion for plant defense suppression. In this study, we uncovered that UPR activity, which is normally restricted to the biotrophic stage *in planta*, inhibits mating and the formation of infectious filaments by Rok1-dependent dephosphorylation of the pheromone responsive MAPK Kpp2. This observation is relevant for understanding how the fungal virulence program is regulated by cellular physiology. UPR-mediated control of mating-type signaling pathways predicts that effector gene expression and the virulence potential are controlled by ER stress levels.

## Introduction

In fungal pathogens, crosstalk between signaling pathways is important for adaptation to the host environment and proper execution of developmental programs (1, 2). The biotrophic plant pathogen *Ustilago maydis* is highly adapted to its host plant *Zea mays* (maize) and plant colonization is prerequisite for completion of its life cycle (3). Pathogenic development is controlled by mating-type signaling pathways coordinating the fusion of two compatible haploid sporidia and formation of the filamentous dikaryon that is capable to infect the plant (4). Plant penetration and establishment of a compatible biotrophic interaction requires rewiring of the mating-type signaling network to adapt to the plant environment and host colonization.

The initial steps of pathogenic development such as cell/cell recognition and fusion of compatible haploid sporidia are controlled by a pheromone (*mfa*)/receptor (*pra*) system encoded by the biallelic *a* mating-type locus (5). Perception of pheromone by the cognate receptor triggers conjugation tube formation, a G2 cell cycle arrest and increased expression of pheromone-responsive genes. Signal transduction within the pheromone response is mediated in parallel by a cAMP-dependent protein kinase A (PKA) and a mitogen-activated protein kinase (MAPK) module to phosphorylate and activate the pheromone response factor 1 (Prf1) (6). Differential phosphorylation of Prf1 by the PKA Adr1 and the MAPK Kpp2 regulates expression of pheromone-responsive genes, including those encoded by the *a* and *b* mating-type loci (7–9).

After cell/cell fusion, all subsequent steps of sexual and pathogenic development are regulated by the heterodimeric bE/bW transcription factor, encoded by the multiallelic *b* mating-type locus. The b-regulated C2H2 zinc finger transcription factor Rbf1 is required and sufficient for all b-dependent processes before plant infection, including filamentous growth, maintenance of the cell cycle arrest and appressoria formation (10). Only after successful plant penetration, the cell cycle arrest is released and proliferation and mitotic division of the dikaryotic filament is initiated. This developmental switch is controlled by the Clp1 protein that is post-transcriptionally regulated and specifically accumulates after plant penetration (11, 12). Clp1 mediates release from the cell cycle block via physical interaction with bW and Rbf1, inhibiting the function of the b-heterodimer and the pheromone response pathway, respectively (12).

The establishment of a compatible biotrophic interaction between *U. maydis* and its host plant maize is mediated by secretion of effector proteins (13, 14). The concerted upregulation of effector gene expression results in a dramatically increased influx into the endoplasmic reticulum (ER), causing ER stress that triggers the unfolded protein response (UPR) as a counter response. The ER membrane-localized stress sensor RNAse/kinase Ire1 promotes expression of the Hac1-like transcriptional activator, termed XBP1 in mammals and Cib1 in *U. maydis*, by endonucleolytic removal of the unconventional intron of the encoding mRNA (15–18). Although the UPR is overall protective by supporting ER homeostasis, prolonged or hyper-activation of the UPR is deleterious for the cell and can lead to UPR-induced cell death (19–21). A functional UPR is critical for ER stress resistance and virulence of pathogenic fungi (22–26). During the life cycle of *U. maydis*, the UPR is specifically activated after plant penetration. Complex formation between the UPR regulator Cib1 and Clp1 initiates fungal proliferation *in planta* by increasing Clp1 stability. By contrast, when activated before plant penetration the UPR inhibits formation of infectious filaments and, as a consequence, virulence in a dose-dependent manner (18). This inhibitory effect is connected to reduced expression of *bE*, *bW* and *rbf1* genes and is independent of Clp1 (18) (for overview see Fig. 1). Hence, additional regulatory connections between the UPR and mating-type pathways in *U. maydis* must exist.

**Figure 1:**
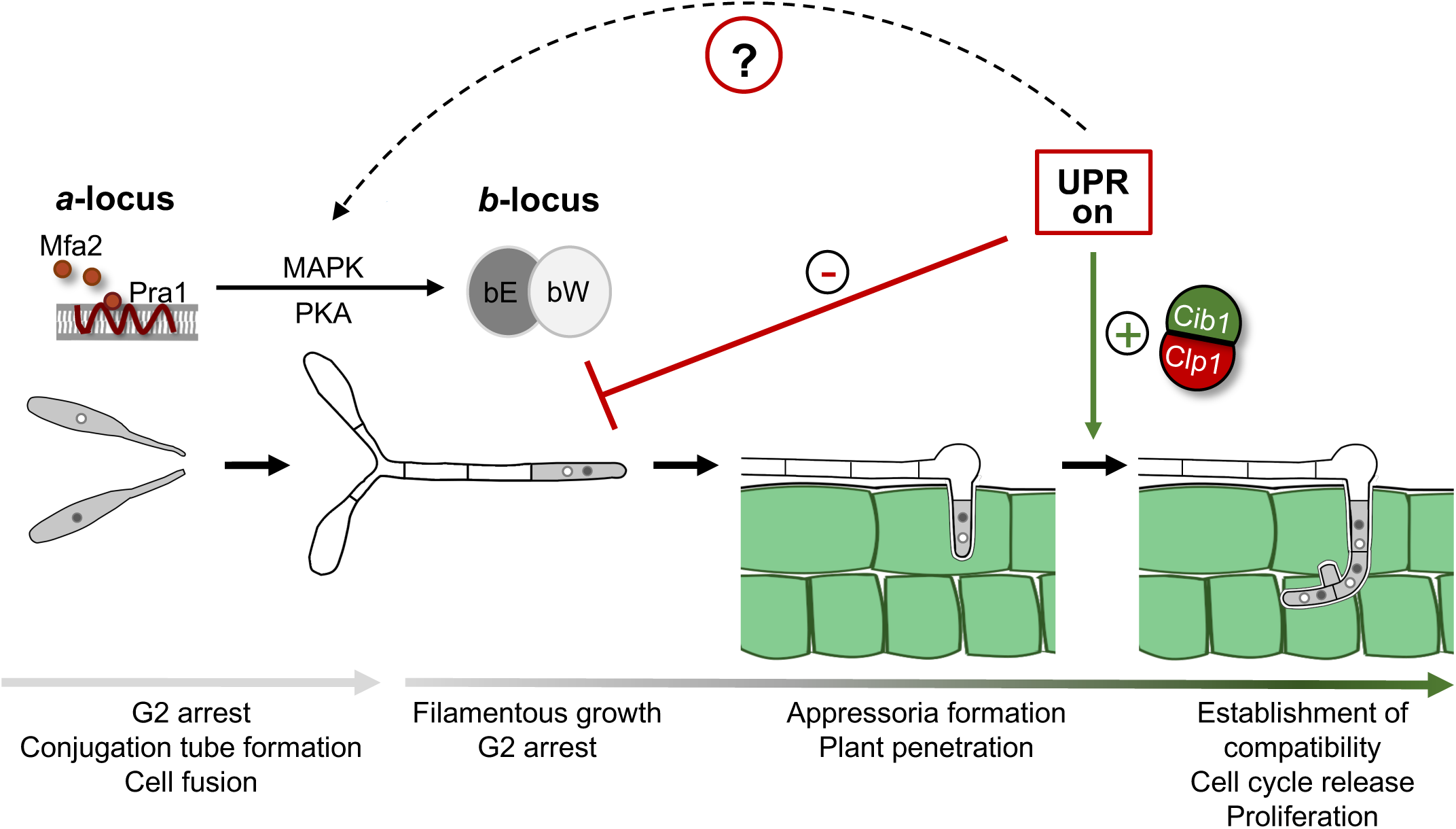
UPR activity is required for fungal proliferation *in planta* but interferes with b-dependent filament formation on the plant surface. Binding of the pheromone (Mfa2) to a compatible the pheromone receptor (Pra1) induces mating of compatible haploid sporidia. The pheromone signal is transmitted by a PKA and a MAPK cascade, inducing a G2 cell cycle arrest and increasing expression of the *a* and *b* mating-type genes (*mfa*/*pra* and *bE*/*bW*). Cell/cell fusion results in formation of the infectious dikaryotic filament. The bE/bW heterodimer controls maintenance of the G2 cell cycle arrest, elongation of the filament and appressoria formation to penetrate the plant surface, After plant penetration, the UPR is activated supporting the release of the cell cycle block by complex formation with Clp1. By contrast, premature UPR activity interferes with b-dependent filament formation and inhibits expression of the *b* mating-type genes. Since this effect is independent of the Clp1 interaction, an active UPR potentially also interacts with mating-type pathways upstream of *bE* and *bW*.

Here we show that an active UPR interferes with the mating-type-dependent signaling upstream of the b-heterodimer. Constitutive activation of the UPR inhibits the morphological and transcriptional response to pheromone. We demonstrate that crosstalk between the UPR and the pheromone regulated MAPK signaling cascade is mediated by Rok1-dependent dephosphorylation of the MAPK Kpp2. Rok1 activity is inversely correlated with fungal virulence and *rok1* deletion fully suppressed the inhibitory effects of the UPR on Kpp2 phosphorylation, filament formation and virulence. Our data suggest that the UPR-controlled activity of mating-type signaling serves as a gain control for fungal virulence that supports ER homeostasis and fungal biotrophy in the *U. maydis*/maize interaction.

## Results

### UPR activity inhibits the *b* mating-type-dependent regulatory network and upstream signaling pathways

To study the effects of a constitutive active UPR on pathogenic development, the intronless *cib1*^s^ mRNA under control of its endogenous promoter was integrated into the *ip* locus of the solopathogenic strain SG200 in single (*cib1*^s^) or multiple copies (*cib1*^s(x)^) (18). In SG200, expression of the compatible *bE1*/*bW2* genes supersedes the need of a mating partner to trigger the pathogenic program und fulfill its life cycle (27). Filament formation of SG200 is suppressed by an active UPR via dose-dependent reduction of *bE*, *bW* and *rbf1* transcript levels (Fig. 2A) (18). Since the b-regulatory network is composed of hierarchically arranged transcription factors, we examined if the complete b-regulatory network is affected by the UPR. To this end, we determined expression levels of the key transcriptional regulators downstream of Rbf1: Biz1, Hdp1 and Hdp2 by quantitative Reverse Transcriptase (qRT)-PCR analysis (10, 28, 29). Consistent with the results obtained for the *b*-genes and *rbf1*, expression levels of all three transcription factors were significantly and dose-dependently reduced by *cib1*^s^ (Fig. 2B). This suggests that UPR activity leads to global suppression of the b-regulatory network.

**Figure 2:**
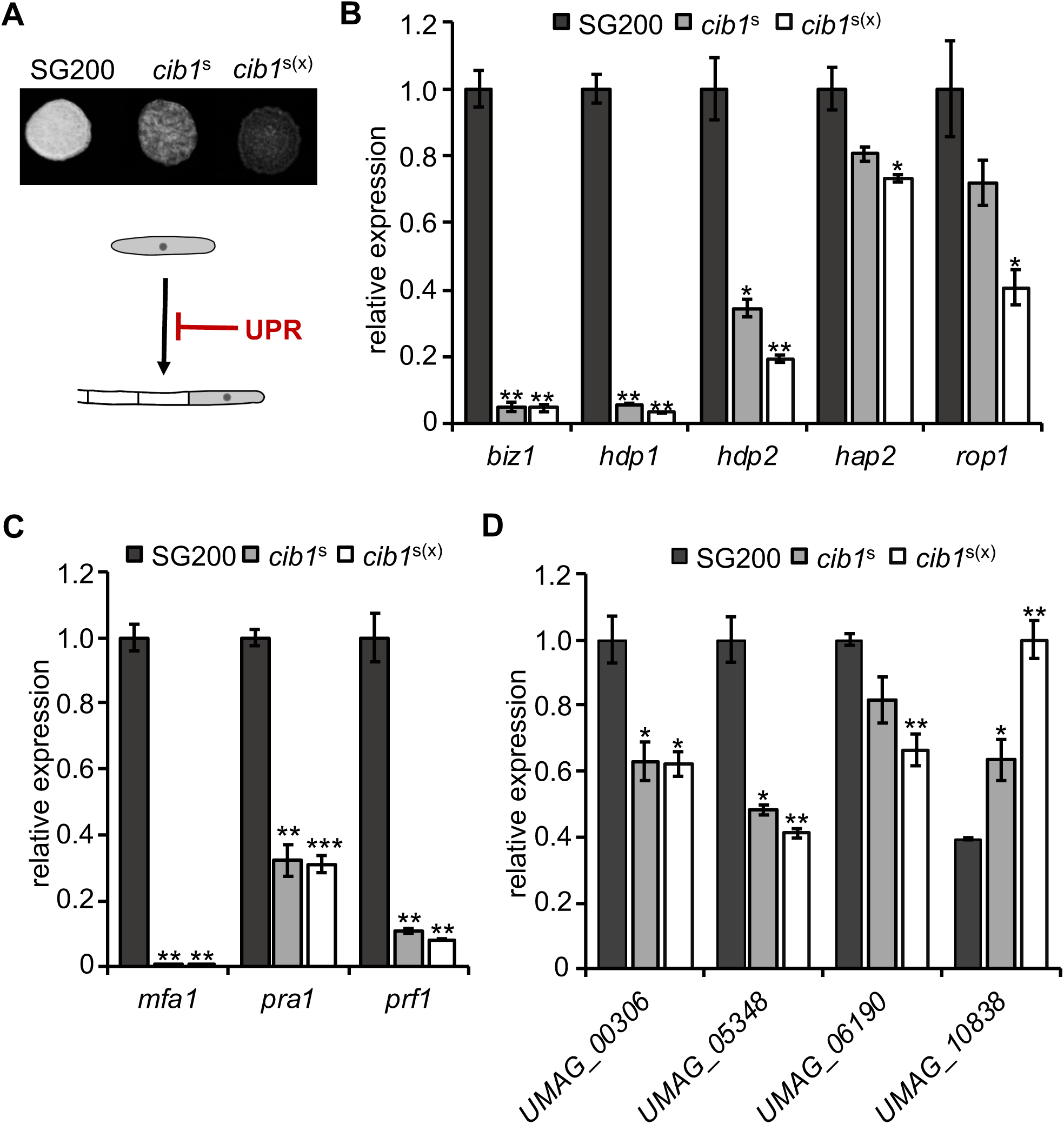
UPR suppresses *b*-dependent filament formation and gene expression. **(A)** Analysis of *b*-dependent filament formation. SG200 and derivatives constitutively expressing *cib1*^s^ or *cib1*^s(x)^ were spotted on potato dextrose solid media supplemented with 1% charcoal. Photos were taken after 24 hours at 28°C. White fuzzy colonies indicate the formation of filaments. **(B-D)** qRT-PCR analysis of gene expression in SG200 and derivatives expressing one (*cib1*^s^) or multiple copies (*cib1*^s(x)^) of *cib1*^s^. *eIF2b* was used for normalization. Expression values represent the mean of three biological replicates with two technical duplicates each. Error bars represent the SEM. Statistical significance was calculated using the student’s *t* test. *P value < 0.05, **P < 0.01, and ***P < 0.001 (B) Expression of *rbf1*-regulated transcription factors *biz1, hdp1* and *hdp2* and regulators of *prf1* expression, *rop1* and *hap2.* (C) Expression of the pheromone genes *mfa1, pra1* and the central transcription factor *prf1.* (D) Expression values of genes *UMAG_00306, UMAG_05348, UMAG_06190* and *UMAG_10838*, which are differentially expressed upon MAPK-dependent phosphorylation of Prf1.

To address at which stage the UPR interferes with the mating-type signaling pathway, we investigated the effects of UPR activity on the pheromone response pathway acting upstream of bE/bW. As a read-out we determined expression levels of the *a* mating-type locus genes *mfa1 and pra1*, encoding the pheromone precursor and the pheromone receptor, respectively, and of *prf1* in SG200 (WT) and the *cib1*^s^-expressing derivative. In comparison to the WT, expression levels of all three genes were significantly reduced in *cib1*^s^-expressing strains (Fig. 2C). Moreover, expression levels of two transcriptional regulators of Prf1, Hap2 and Rop1, and of the Prf1-induced target genes *UMAG_00306, UMAG_05348* and *UMAG_*06190 (8) were suppressed by the UPR as well (Fig. 2D). Interestingly, UPR activity also affected Prf1-dependent repression of *UMAG_10838* (Fig. 2D). This suggests that the UPR interferes with mating-type signaling upstream of the b-heterodimer by negatively affecting Prf1-dependent gene regulation.

### UPR activity impedes the morphological and transcriptional responses to pheromone

In the solopathogenic strain SG200, expression of a compatible pheromone/pheromone receptor pair and compatible *b*-genes leads to constitutive activation of the *a* and *b* mating-type pathways. Both pathways are connected by various feedback loops (7, 8, 12), complicating their functional analysis. We thus used the haploid FB1 and FB2 WT strains to specifically address the impact of the UPR on the pheromone response (controlled by the *a* mating-type locus). Similar to the results obtained in the SG200 background, mating assays between compatible FB1 and FB2 strains revealed reduced filament formation in crossings between *cib1*^s^-expressing derivatives on charcoal containing solid media (Fig. S1). When liquid cultures of FB1 (WT) and the *cib1*^s^-expressing derivative were treated with synthetic *a2* pheromone, conjugation tube formation was inhibited in *cib1*^s^-expressing strains compared to the WT (Fig. 3A). Conjugation tube formation was significantly reduced from 32.9% (WT) to 17.3% in FB1*cib1*^s^ and to 4.5% in FB1*cib1*^s(x)^ strains (Fig. 3B). This demonstrates that the UPR inhibits the morphological response to pheromone in a dose-dependent manner.

**Figure 3:**
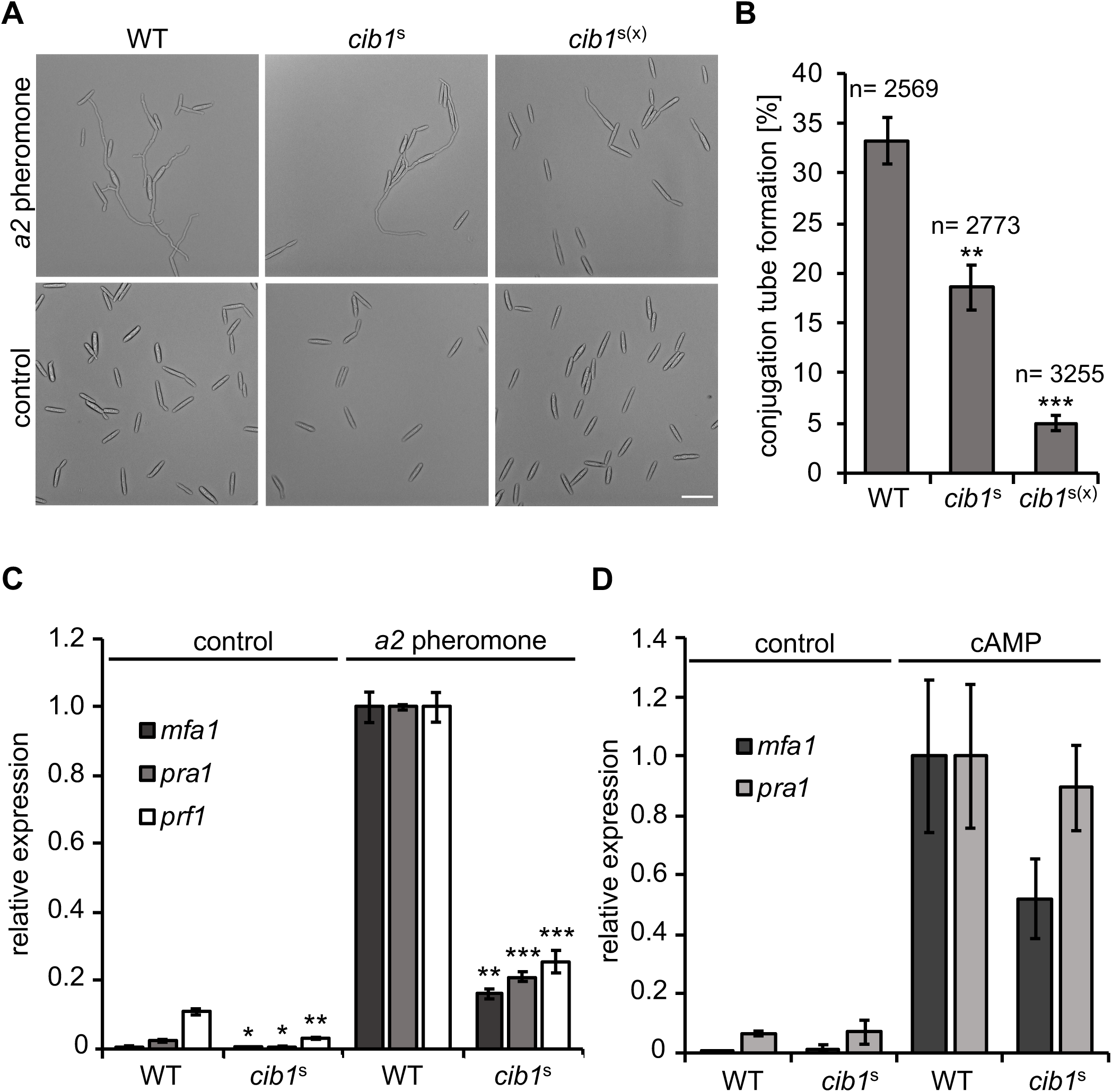
The morphological and transcriptional response to pheromone is inhibited by UPR activity. **(A)** Microscopic analysis of pheromone-induced conjugation tube formation. Wild type strain FB1 (WT) and derivatives constitutively expressing *cib1*^s^ or *cib1*^s(x)^ were treated with synthetic *a2* pheromone or DMSO as negative control and grown for 6 hours at 28°C under rotation. Formation of conjugation tubes was monitored by bright field microscopy. Bar = 10 µm. **(B)** Quantification of conjugation tube formation. Depicted values represent mean of three independent experiments. Error bars represent the SD. n = number of total cells counted. Statistical significance was calculated using the student’s *t* test. **P value < 0.01, and ***P < 0.001. **(C)** qRT-PCR analysis of *mfa1, pra1* and *prf1* gene expression in FB1 (WT) and derivatives expressing *cib1^s^* after treatment with synthetic *a2* pheromone. Cells were treated with synthetic *a2* pheromone (2.5 µg/ml) for 6 hours at 28°C. *eIF2b* was used for normalization. Expression values represent the mean of three biological replicates with two technical duplicates each. Error bars represent the SEM. Statistical significance was calculated using the students *t* test. *P value < 0.05, **P < 0.01, and ***P < 0.001. **(D)** qRT-PCR analysis of genes *mfa1* and *pra1* in FB1 (WT) and derivatives expressing *cib1^s^* after treatment with cAMP. Cells were treated with 6 mM cAMP for 12 hours at 28°C. *eIF2b* was used for normalization. Expression values represent the mean of two biological replicates with two technical duplicates each. Error bars represent the SEM.

We next tested if suppression of the pheromone-induced morphological response correlates with an altered transcriptional response. qRT-PCR analysis revealed that basal and pheromone-induced expression levels of *mfa1*, *pra1* and *prf1* were significantly lower in the *cib1*^s^-expressing strain compared to WT (Fig. 3C). This suggests that the UPR inhibits both, the morphological and the transcriptional response to pheromone.

Since signal transduction within the pheromone response is mediated by PKA- and MAPK-pathways, we tested whether UPR activation affects the activity of the cAMP-dependent PKA pathway. To specifically activate the PKA pathway, we treated FB1 (WT) and FB1*cib1^s^* cells with cAMP (6 mM) and monitored expression of *mfa1* and *pra1* genes. This revealed that cAMP-induced expression of *mfa1* and *pra1* was slightly lower in *cib1*^s^-expressing strains when compared to FB1 (Fig. 3D). However, under these conditions the inhibitory effect of an active UPR on *mfa1* and *pra1* expression was modest and less pronounced when compared to pheromone treated cells. Therefore, the strong impact of an active UPR on the pheromone response pathway is most likely not achieved via the cAMP-dependent PKA signaling pathway.

### Genetic activation of the pheromone responsive MAPK pathway reveals differential inhibition of morphological and transcriptional responses by the UPR

To address a potential interaction between the UPR and the MAPK pathway, we genetically activated the pathway by expression of the active MAPK kinase (MAPKK) *fuz7^DD^* in FB1 (WT) and FB1*cib1*^s^ strains (6, 30). Expression of Fuz7^DD^ by the arabinose-inducible *crg1*- promoter results in formation of filamentous cells, resembling conjugation tubes (6). Under these conditions, 86.8% of the WT cells and 95.9% of *cib1*^s^-expressing cells reacted with the formation of conjugation tube-like structures (Fig. 4A and B). Conversely, transcript levels of the pheromone genes *mfa1*, *pra1* and of *prf1* were significantly less induced by Fuz7^DD^ in *cib1*^s^-expressing cells compared to the WT (Fig. 4C). Hence, the UPR inhibits the Fuz7^DD^-induced gene regulation but not the morphological response. Importantly, transcriptional regulation of the *mfa1* and *pra1* genes is Prf1-dependent, whereas the morphological response is not (6, 31). This indicates that the UPR negatively affects Prf1 activity, which is regulated by phosphorylation of the MAPK Kpp2.

**Figure 4:**
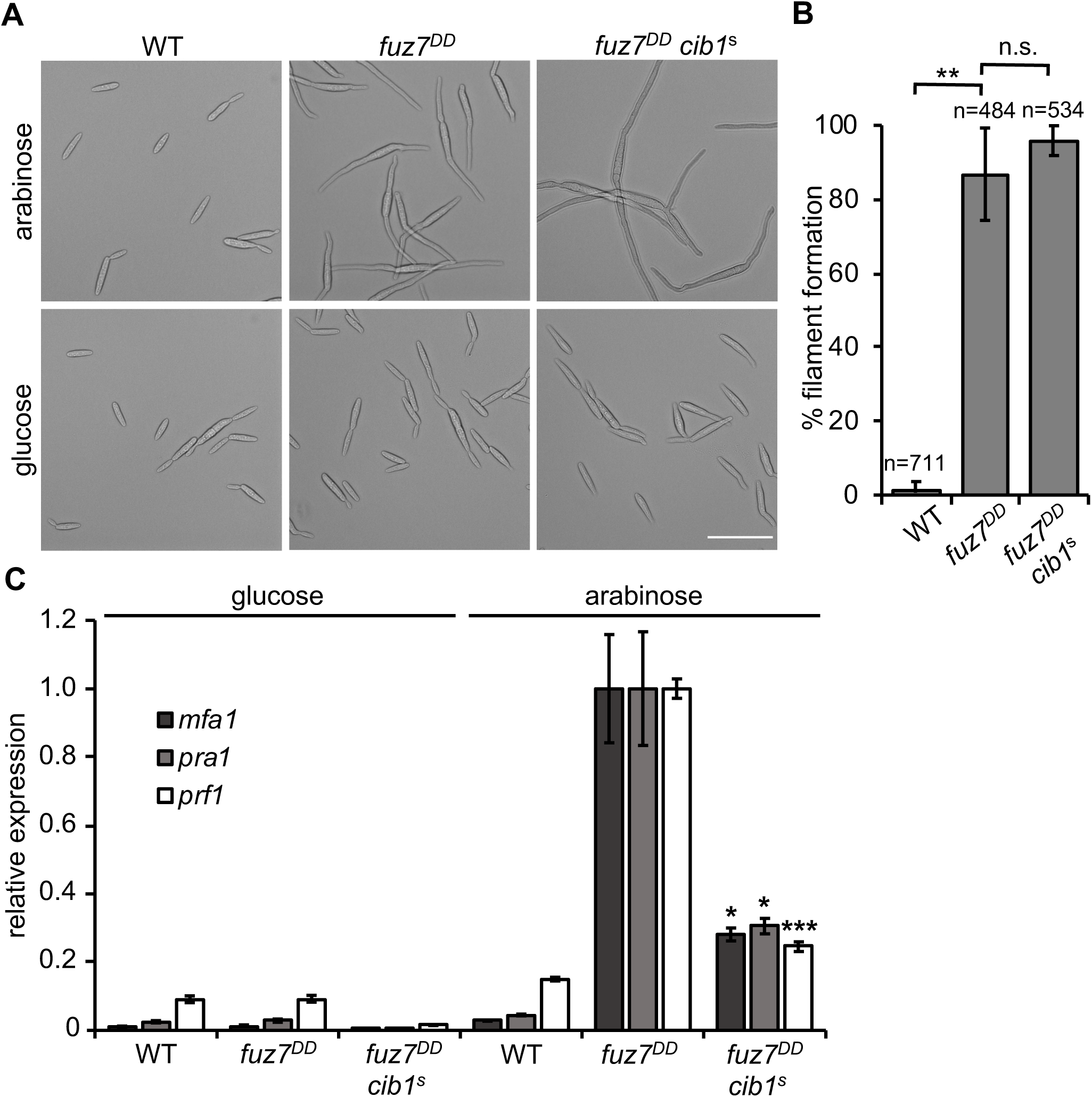
The UPR differentially affects Fuz7^DD^-induced morphological and transcriptional responses. **(A)** Microscopic analysis of *fuz7^DD^*-induced conjugation tube-like structures. Strain FB1 (WT), FB1 *fuz7^DD^* (*fuz7^DD^* expression under control of the arabinose-inducible *crg1* promoter) and a derivative expressing *cib1^s^* were grown for 6 hours at 28°C in CM-glucose (non-inducing) or CM-arabinose (inducing) medium. Formation of conjugation tube-like structures was monitored by bright field microscopy. Bar = 10 µm. **(B)** Quantification of formation of conjugation tube-like structures. The bars represent mean value of three independent experiments. Error bars represent the SD. n = number of total cells counted. Statistical significance was calculated using the students *t* test. **P value < 0.01. **(C)** qRT-PCR analysis of *mfa1, pra1* and *prf1* gene expression in FB1 (WT), FB1 *fuz7^DD^* (*fuz7^DD^* expression under control of the arabinose-inducible *crg1* promoter) and a derivative expressing *cib1^s^*. Cells were grown for 6 hours at 28°C in CM-glucose (non-inducing) or CM-arabinose (inducing) medium. *eIF2b* was used for normalization. Expression values represent the mean of three biological replicates with two technical duplicates each. Error bars represent the SEM. Statistical significance was calculated using the students *t* test. *P value < 0.05 and ***P < 0.001.

### Phosphorylation of the MAPK Kpp2 but not the interaction with the MAPKK Fuz7 is suppressed by UPR-activity

Since the MAPK Kpp2 controls the morphological response to pheromone and Prf1 activity (32–34), we reasoned that the UPR might affect the MAPK cascade at the level of Kpp2. To test our hypothesis, we expressed a functional Kpp2-GFP fusion protein under the control of its endogenous promoter from the endogenous genomic locus in WT (FB1*fuz7^DD^*) and *cib1*^s^-expressing (FB1*fuz7^DD^ cib1*^s^) strains. Western hybridization revealed that abundance of Kpp2-GFP was not altered by induced expression of *fuz7^DD^* or *cib1*^s^-mediated UPR activity (Fig. 5A). By contrast, phosphorylation of the Kpp2-GFP fusion protein was strongly increased upon induced expression of *fuz7^DD^* in WT but remained almost undetectable in *cib1*^s^-expressing strains under identical conditions (Fig. 5A). This suggests that an active UPR negatively affects phosphorylation of Kpp2.

**Figure 5:**
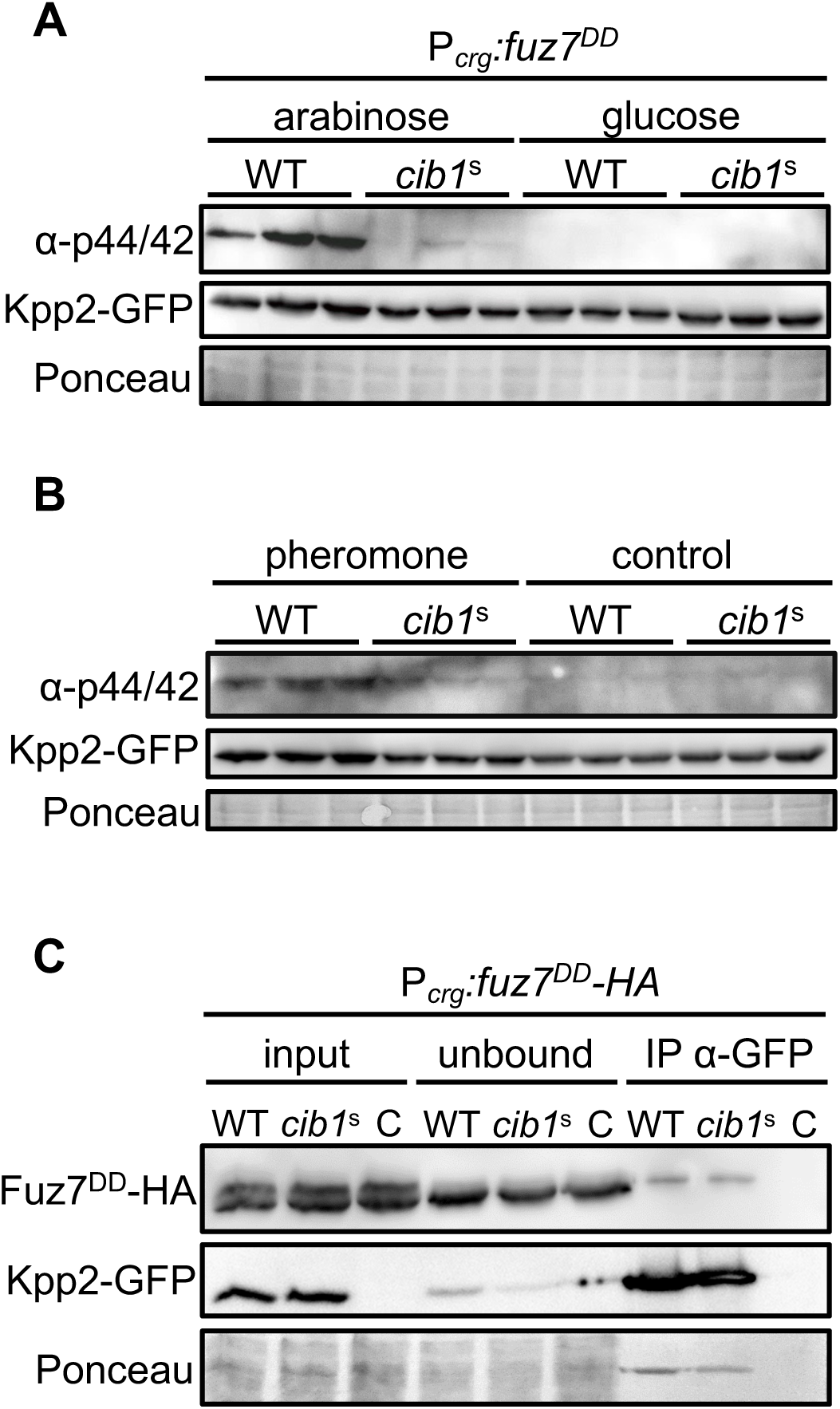
Expression of *cib1^s^* diminishes Kpp2 phosphorylation but not the interaction to Fuz7. **(A)** Immunoblot analysis of Kpp2-GFP protein levels and phosphorylation. Derivatives of strains FB1 *fuz7^DD^* (*fuz7^DD^* expression under control of the arabinose-inducible *crg1* promoter, WT) and FB1 *fuz7^DD^ cib1^s^* (*cib1^s^*) expressing a Kpp2-GFP fusion protein were grown in CM-glucose (non-inducing) or CM-arabinose (inducing) liquid medium for 6 hours at 28°C. Kpp2-GFP was visualized using a GFP-specific antibody. Phosphorylation of Kpp2-GFP was visualized using the α-phospho44/42 antibody that specifically recognizes the phosphorylated TEY motif of MAPKs. The Ponceau stained membrane served as loading control. **(B)** Immunoblot analysis with the same strains as in (A) treated with synthetic *a2* pheromone for 6 hours at 28°C. The same antibodies as in (A) were used to visualize Kpp2-GFP and MAPK phosphorylation. The Ponceau S stained membrane served as loading control. **(C)** Co-immunoprecipitation analysis of Kpp2-GFP and Fuz7^DD^-HA. Derivatives of strains FB1 *fuz7^DD^ kpp2-GFP* (WT) and FB1 *fuz7^DD^ kpp2-GFP cib1*^s^ (*cib1*^s^) containing a *fuz7^DD^-HA* fusion protein under control of the arabinose-inducible *crg1* were grown in CM-arabinose (inducing) liquid medium for 6 hours at 28°C. A strain expressing untagged Kpp2 served as negative control (C). Total protein extract was subjected to co-immunoprecipitation using GFP-specific beads. Fuz7^DD^-HA was visualized using an HA-specific antibody. Kpp2-GFP was visualized using an GFP-specific antibody. The Ponceau stained membrane served as loading control.

Fuz7^DD^-mediated genetic activation of the MAPK pathway relies on an artificial expression system that bypasses feedback regulation within the MAPK pathway. To address UPR-dependent effects in a more “natural” setting, we monitored Kpp2 phosphorylation in response to pheromone treatment in FB1 (WT) and *cib1*^s^-expressing strains. Consistently, expression of *cib1*^s^ did not affect the abundance but specifically reduced phosphorylation levels of Kpp2-GFP in response to pheromone (Fig. 5B), confirming our previous results obtained upon genetic activation of the MAPK pathway (Fig. 5A).

Signal transduction within MAPK cascades requires the physical interaction between pathway components that is supported by accessory or scaffold proteins to establish signaling modules and foster phospho-transfer between MAPK-pathway components (35, 36). To address if the UPR affects the interaction between Fuz7 and Kpp2, we expressed functional Kpp2-GFP and Fuz7^DD^-HA fusion proteins in WT (FB1) and *cib1*^s^-expressing strains and immunoprecipitated Kpp2-GFP by GFP-trap. This revealed that Fuz7^DD^-HA was co-immunoprecipitated to similar amounts in the WT and *cib1*^s^-expressing strain. By contrast, Fuz7^DD^-HA was not co-immunoprecipitated in the control strain expressing untagged Kpp2 (Fig. 5C). This indicates that the Kpp2/Fuz7 interaction is specific and not affected by an active UPR.

### Deletion of *rok1* restores Kpp2 phosphorylation, filament formation and virulence of *cib1*^s^-expressing strains

Phosphorylation levels of MAPKs are determined by relative activities of phosphorylating and dephosphorylating proteins (37, 38). Previously, it has been shown that the dual specificity phosphatase (DSP) Rok1 mediates dephosphorylation of Kpp2 and thereby counteracts activity of the pheromone response pathway (39). To test if the inhibitory effect of an active UPR is mediated by Rok1, we deleted *rok1* in the WT (FB1P*_crg_*:*fuz7^DD^*) and the *cib1*^s^-expressing derivative (FB1P*_crg_*:*fuz7^DD^* P*_cib1_:cib1*^s^). Consistent with our previous results, Kpp2-GFP abundance was not altered under any of the tested conditions. Strikingly, induced expression of *fuz7^DD^* resulted in Kpp2-GFP phosphorylation in WT and *cib1*^s^-expressing Δ*rok1* cells to similar levels (Fig. 6A). Hence, deletion of *rok1* fully repressed the negative effect of an active UPR on Kpp2 phosphorylation.

**Figure 6:**
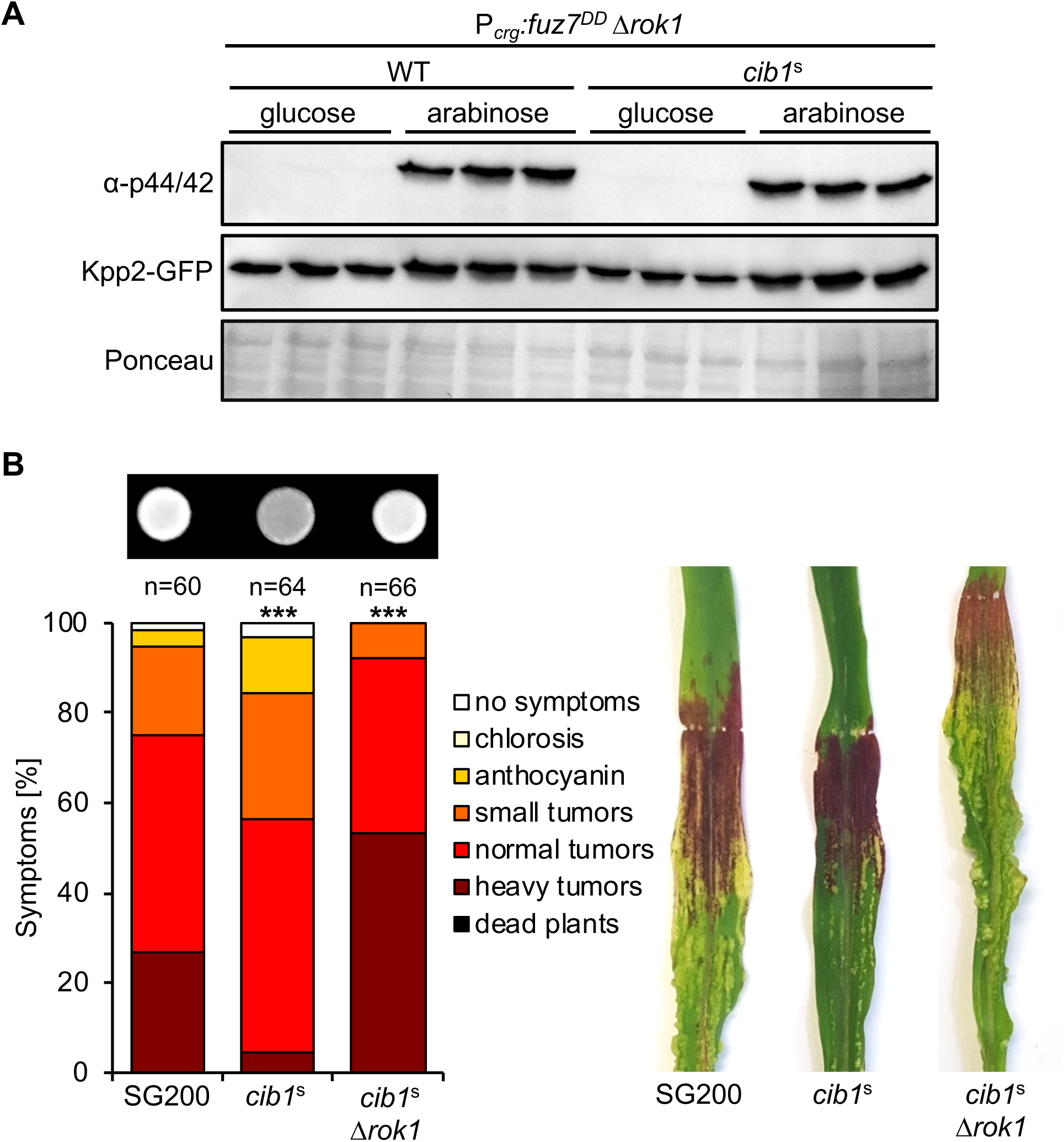
Deletion of *rok1* suppresses *cib1*^s^-dependent inhibition of Kpp2 phosphorylation, filament formation and virulence. **(A)** Immunoblot analysis of Kpp2-GFP protein levels and phosphorylation in *rok1* deletion strains. Derivatives of strains FB1 *fuz7^DD^* (WT) and FB1 *fuz7^DD^ cib1*^s^ (*cib1*^s^) deleted for *rok1* were grown in CM-glucose (non-inducing) or CM-arabinose (inducing) liquid medium for 6 hours at 28°C. Protein extracts were subjected to immunoblot analysis, Kpp2-GFP levels were visualized using a GFP-specific antibody. Phosphorylation of Kpp2-GFP was visualized with an α-phospho44/42 antibody specifically recognizing the phosphorylated TEY motif of MAPKs. The Ponceau S stained membrane served as loading control. **(B)** Analysis of *b*-dependent filament formation and virulence of *rok1* deletion strains. SG200, SG200*cib1*^s^ and a derivative deleted for *rok1* (*cib1*^s^Δ*rok1*) were spotted on potato dextrose solid media supplemented with 1% charcoal. Photos were taken after 24 hours at 28°C. White fuzzy colonies indicate the formation of filaments. For plant infection assay, strains were inoculated into 8-day-old maize seedlings. Disease symptoms were rated 8 days after inoculation (dpi) and grouped into categories as shown in the figure legend. n = number of inoculated plants. Photos were taken at 12 dpi and represent the most common infection symptom. Statistical significance was calculated using the Mann-Whitney-test. ***P value< 0.001.

We previously demonstrated that constitutive UPR activity inhibits filament formation and virulence of the solopathogenic strain SG200 (18). These effects are reminiscent of the phenotype observed upon constitutive overexpression of *rok1* (39). By contrast, *rok1* deletion mutants show increased filament formation and are hypervirulent in plant infection assays (39). To test if deletion of *rok1* not only restores phosphorylation of Kpp2 in *cib1*^s^-expressing strains, but also filament formation and virulence, we deleted *rok1* in the SG200*cib1*^s^ strain. Strikingly, both, reduced filament formation and virulence of strain SG200*cib1*^s^ was fully restored in the Δ*rok1* derivative (Fig. 6B). Interestingly, virulence of SG200*cib1*^s^ *rok1* was not only restored to WT levels, but was further increased in comparison to the WT control resembling the hypervirulent phenotype of SG200Δ*rok1* (39). Conclusively, our data show that UPR-dependent inhibition of mating-type signaling is mediated via the DSP Rok1.

### UPR activity does not affect Rok1 transcript or protein levels

To investigate how the UPR regulates Rok1, we determined Rok1 transcript and protein levels in the WT (FB1P*_crg_*:*fuz7^DD^*) and the *cib1*^s^-expressing derivative. This revealed that *rok1* transcript levels were only slightly reduced in *cib1*^s^-expressing strains compared to the WT control under both, *fuz7^DD^*-repressing (glucose) and -inducing (arabinose) conditions (Fig. S2A). To test for potential UPR-induced alterations of Rok1 protein levels, we expressed a Rok1-mCherry fusion protein under control of the endogenous promoter from the endogenous genomic locus in WT and *cib1*^s^-expressing strains. Since the fusion protein was not detectable by Western hybridization, Rok1-mCherry was enriched by immunoprecipitation. Total protein amounts of *U. maydis* protein extracts were adjusted and immunoprecipitation was performed with an increased amount of mCherry-Trap beads (Chromotek) to prevent saturation of the beads and enable a quantitative comparison between both strains. While Rok1-mCherry was not detectable under non-induced conditions, protein levels were similar in both strains in response to *fuz7^DD^*-mediated genetic activation of the MAPK pathway (Fig. S2B). Taken together, these results suggest that the UPR regulates Rok1 on a posttranslational level, potentially by influencing Rok1 activity.

### The MAPK Kpp2 is dispensable for biotrophic development *in planta*

During the life cycle of *U. maydis*, the UPR is specifically activated after plant penetration (18). Hence, the crosstalk between the UPR and the MAPK pathway is supposedly involved in controlling biotrophic growth *in planta*. Kpp2 is important for the mating process and appressoria formation (32), whereas the partially redundant MAPK Kpp6 is expressed in a bE/bW-dependent manner and its phosphorylation is required for penetration of the plant surface (40).

To address the potential function of Kpp2 after plant penetration, we used a conditional gene expression approach, allowing the stage-specific depletion of gene expression during development *in planta*. FB1Δ*kpp2* and FB2Δ*kpp2* strains were transformed with constructs expressing *kpp2* under control of the conditional promoters P*_UMAG_12184_* and P*_UMAG_03597_*. These promoters are active during the first two (P*_UMAG_12184_*) or four (P*_UMAG_03597_*) days after inoculation (the detailed description of our conditional gene expression approach will be published elsewhere) but show strongly reduced activity at subsequent stages of pathogenic growth *in planta* (41). We used compatible mixtures of FB1 and FB2 (WT) and derivatives thereof for plant infection experiments. Consistent with previous studies, virulence of compatible combinations of Δ*kpp2* strains was significantly impaired when compared to the WT (Fig. 7) (32). Importantly, when *kpp2* was expressed under control of P*_UMAG_12184_* or P*_UMAG_03597_* promoters in the Δ*kpp2* background (FB1Δ*kpp2* P*_UMAG_12184_:kpp2* x FB2Δ*kpp2* P*_UMAG_12184_:kpp2*; FB1Δ*kpp2* P*_UMAG_03597_:kpp2* x FB2Δ*kpp2* P*_UMAG_03597_:kpp2*), virulence was fully restored (Fig. 7). Hence, *kpp2* is required for the mating process and important for appressoria formation but dispensable for pathogenic development *in planta.* As we observed that Rok1 interacts with Kpp2 and Kpp6 in the yeast two-hybrid system (Fig. S3) and that both MAP kinases are dephosphorylated by Rok1 (42), it is conceivable that the crosstalk between UPR and the mating-type pathways might as well affect the function of both proteins (40).

**Figure 7:**
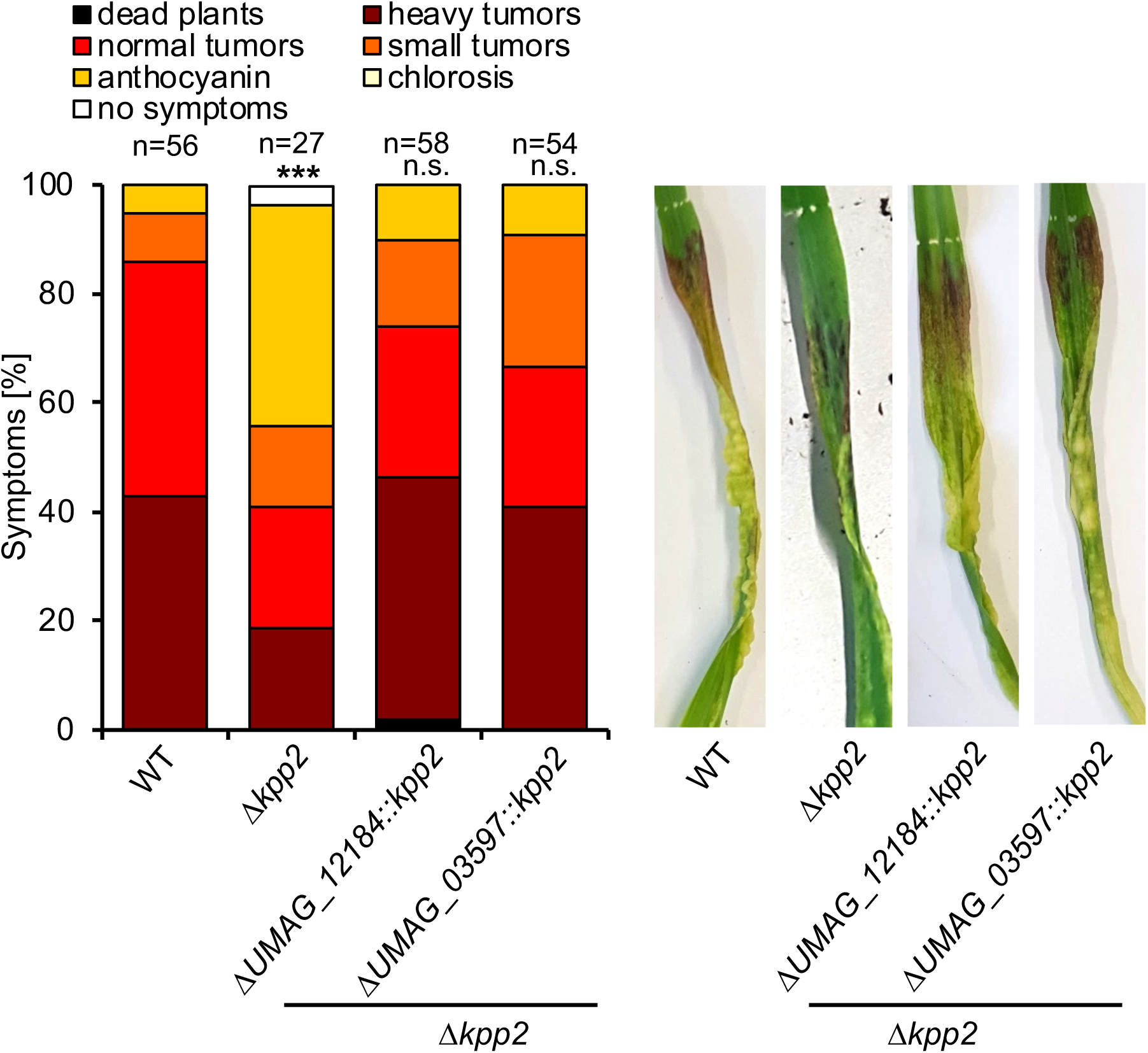
Kpp2 is dispensable for biotrophic development *in planta*. Plant infection assay with compatible mixtures of the haploid strains FB1 x FB2 (WT), FB1*Δkpp2* x FB2*Δkpp2* (*Δkpp2*) and derivatives of *Δkpp2* strains expressing P*_UMAG_12184_:kpp2* or P*_UMAG_03597_:kpp2* were inoculated into 8-day-old maize seedlings. Disease symptoms were rated 8 days after inoculation and grouped into categories as shown in the figure legend. n = number of inoculated plants. Two independent clones were investigated for conditional *kpp2* expression. Photos were taken at 8 dpi and represent the most common infection symptom. Statistical significance was calculated using the Mann-Whitney-test. ***P value < 0.001.

**Figure 8:**
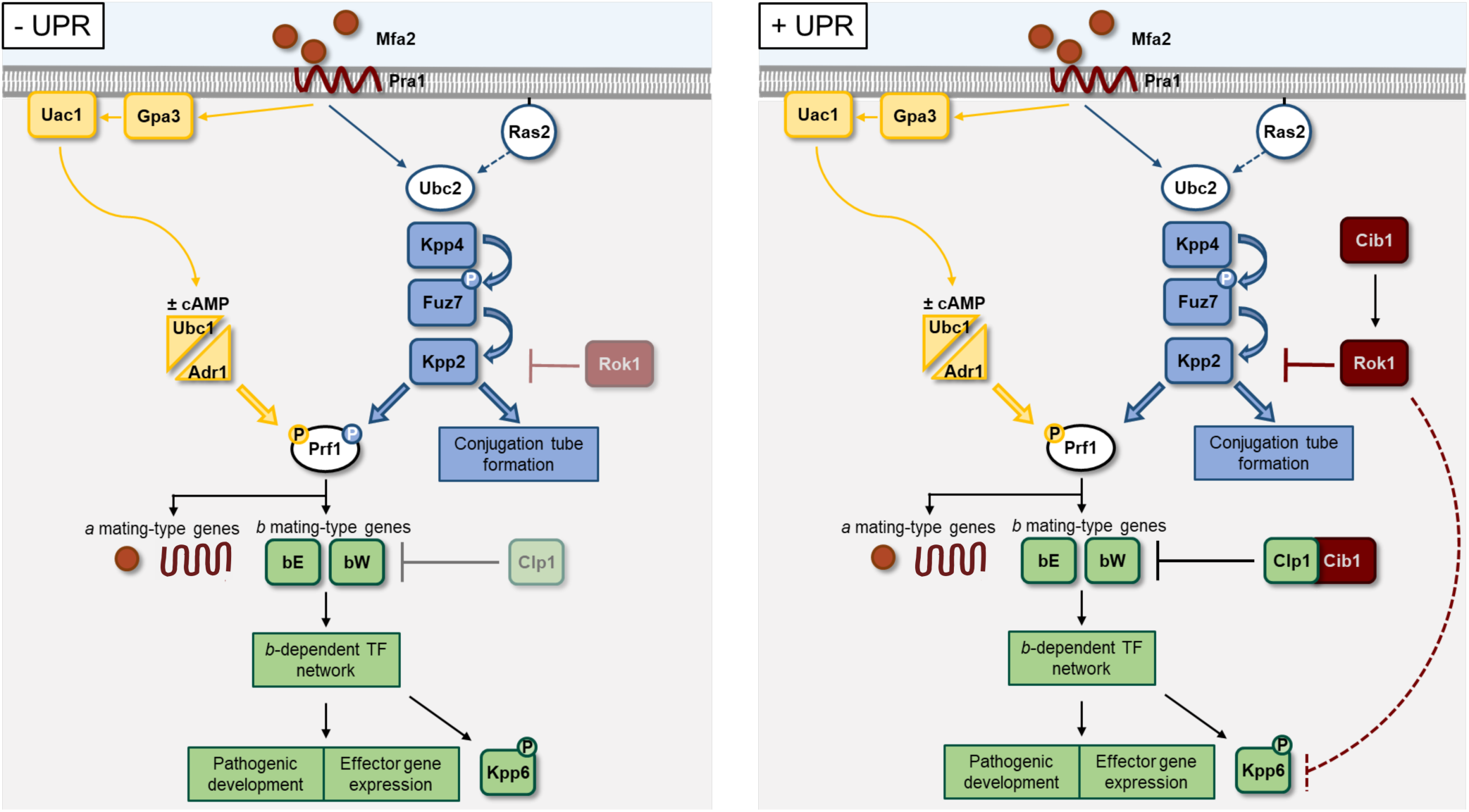
Proposed model of crosstalk between the UPR and mating-type pathways. Binding of pheromone (Mfa2) to the cognate pheromone receptor (Pra1) activates MAPK (blue) and a PKA (yellow) signaling cascades resulting in phosphorylation and activation of the transcriptional regulator Prf1. Active Prf1 induces expression of the *a* and *b* mating-type genes, while the terminal MAPK Kpp2 controls formation of conjugation tubes and mating. After cell/cell fusion, the *b*- heterodimer-regulated transcription factor network controls further pathogenic development and expression of effector genes. After penetration of the plant surface, the UPR (dark red) is activated (+UPR), promoting fungal proliferation through Cib1/Clp1 complex formation and establishment of the compatible biotrophic interaction via efficient effector secretion. Crosstalk of the UPR and the DSP Rok1 under increased ER stress constitutes a negative feedback on *a*- and *b*-dependent signaling pathways by (partially) inactivating Kpp2 (and potentially Kpp6), ultimately reducing effector gene expression and thereby ER-stress levels.

## Discussion

In *U. maydis*, the UPR is connected to the control of biotrophic growth via the interaction between the Hac1-like UPR regulator Cib1 and Clp1. In this study, we show that the UPR affects pathogenic development of *U. maydis* on additional levels. While the plant-specific activation of the UPR fosters pathogenic growth *in planta*, premature UPR activation counteracts pathogenic development prior host plant penetration. Hence, unfit or stressed cells are less likely to undergo mating than non-stressed cells. We observed that UPR activity counteracts pheromone-dependent signaling by suppression of Kpp2 phosphorylation in a *rok1*-dependent manner, identifying Rok1 as a novel regulatory target of the UPR.

Gene expression analysis confirmed the global inhibitory effect of the UPR on the *b-* dependent transcription factor network (18). Moreover, constitutive UPR signaling suppressed the morphological and transcriptional response to pheromone. By contrast, in the human pathogenic fungus *Cryptococcus neoformans*, deletion of *IRE1* resulted in increased pheromone gene expression, but defects in sexual mating (43). Sensing of compatible pheromone by haploid *U. maydis* cells results in activation of the cAMP-dependent PKA Adr1 and MAPK-mediated signaling pathway, finally leading to differential phosphorylation and activation of Prf1 by Adr1 and Kpp2. Kpp2 itself is activated by phosphorylation and regulates the transcriptional pheromone response via Prf1 and the morphological response in a Prf1-independent manner (6). After pheromone treatment, both responses are inhibited by an active UPR (Fig. 3), whereas only pheromone gene expression but not the formation of conjugation tubes was suppressed after genetic activation of the pheromone pathway. These findings are in line with previous results, illustrating that the morphological and the transcriptional response to pheromone bifurcate downstream of Kpp2 (6). Importantly, phosphorylation of Kpp2 was strongly reduced by the UPR under these conditions. Since strains expressing non-phosphorylatable Kpp2^AEF^ fail to form conjugation tubes upon pheromone treatment (6), our results imply that low levels of Kpp2 phosphorylation might be sufficient for conjugation tube formation but not for Kpp2-dependent Prf1 activation.

The interaction between the UPR and MAPK pathways is not limited to *U. maydis*. In the budding yeast *Saccharomyces cerevisiae*, nitrogen-induced pseudohyphal growth, which depends on MAPK and PKA pathways is inhibited by an active UPR (44). However, a direct connection between the UPR and phosphorylation of a pheromone-dependent MAPK has not been reported. In the human pathogens *Aspergillus fumigatus* and *C. neoformans*, a functional UPR is required for cell wall stress resistance, suggesting connections to the cell wall integrity (CWI) pathway (45–47). In budding yeast, activation of the CWI pathway triggers the UPR (48) and the UPR is augmented by delayed PKA signaling, leading to an extended secondary response to ER stress (49). These connections do not appear to be conserved in *U. maydis*: deletion of *cib1* did not affect resistance to cell wall perturbing agents in *U. maydis* and UPR activity did not substantially affect expression of PKA subunits, the PKA-repressed regulator Hgl1 (50) or the PKA-dependent induction of *mfa1* and *pra1* genes.

To allow tight control of signaling, MAP kinase phosphatases (MKP) are regulated on multiple levels, including transcriptional and post-translational control (51). Transcription of *MSG5*, the *rok1* ortholog in *S. cerevisiae*, is induced by pheromone stimulation, placing Msg5p in a negative feedback loop with the pheromone signaling cascade (52). Similarly, *rok1* transcript levels in *U. maydis* were elevated upon activation of the MAPK module (39). However, since neither transcript nor Rok1 protein levels were altered by an active UPR, additional, post-translational mechanisms must exist regulating Rok1 activity under these conditions. Post-translational modifications of MKPs like phosphorylation and ubiquitination can alter subcellular localization, stability and specificity of phosphatases. *S. cerevisiae* Msg5p is phosphorylated by its MAPK targets Fus3 and Slt2 as part of a regulatory feedback mechanism (52, 53). In the hemibiotrophic blast fungus *Magnaporthe oryzae* (*Pyricularia oryzae*), the MKP Pmp1 dephosphorylates the MAPKs Pmk1 (Fus3/Kpp2 ortholog) and Mps1 upon phosphorylation at a conserved serine residue (54). Accordingly, two potential phosphorylation sites have been predicted for Rok1 (55), but their functionality has not been addressed, yet.

Deletion of *rok1* results in hypervirulence of *U. maydis* and restored phosphorylation of Kpp2, filament formation and virulence in strains with active UPR, indicating a UPR-dependent regulation of Rok1. Previous studies suggested that hypervirulence of *rok1* deletion mutants is connected to increased activity of the pheromone response pathway and thus elevated expression of *bE* and *bW* genes (39). Consistently, increased virulence was also observed in strains overexpressing the pheromone receptor Pra2 (FB1 P*_otef_:pra2*) (56), even allowing to partially bypass the loss of the MAPK Crk1 that is crucial for full activity of the signaling pathway (57). This suggests that activity of the mating-type pathways correlates with aggressiveness of *U. maydis*. As a biotrophic pathogen, successful completion of its life cycle is dependent on the living host plant. In this way, the UPR-dependently increased activity of Rok1 might support the coordinated and well-balanced growth of *U. maydis* inside the plant ensuring host survival during fungal colonization.

Another important role of this regulatory connection for biotrophic growth *in planta* is predicted based on the influence on the b-regulatory network. Expression of effector-encoding genes during pathogenic development is mainly controlled by the b-heterodimer or *b*-regulated transcription factors, including Rbf1, Hdp2 and potentially Biz1 (10, 28, 29, 41, 58). Their proper folding, processing and secretion, however, is dependent on a functional UPR (50, 59, 60). Deleterious hyperactivation of the UPR is suppressed by Clp1-dependent modulation of UPR gene expression (18, 50). In parallel, our data predict that increased ER stress levels initiate a Rok1-mediated negative feedback on these regulators. Consequently, effector gene expression is temporarily reduced to support recovery from excessive ER stress and achieve ER homeostasis. Importantly, transcription of the Pit2 effector, which mediates suppression of host cysteine proteases (61, 62), is directly regulated by Cib1 and, in addition, by Hdp2 (29, 60). As a consequence, *pit2* expression is likely not to be affected by UPR-dependent inhibition of the mating-type regulatory network, which is in line with the pivotal function of Pit2 in *U. maydis* virulence (63).

Kpp2 is highly similar to the unconventional MAPK Kpp6 (40) and both MAPKs are targeted for dephosphorylation by Rok1 (42). While Kpp2 is important for mating, Kpp6 is not expressed or required at this stage. However, both MAPKs are partially redundant during appressoria formation (40). While the single deletion of either MAPK resulted in reduced pathogenicity, only the double deletion completely abolished virulence of *U. maydis* (39, 40). Conditional gene expression revealed that Kpp2 is not required for later stages of biotrophic development *in planta*, suggesting that Kpp6 might be redundant to Kpp2 also at this stage. The physical interaction between Rok1 and both MAP kinases implies a similar and direct mode of action and suggests that the UPR affects phosphorylation levels of Kpp6 as well. Hence, we cannot exclude that Kpp6 is the functionally important Rok1 target with respect to biotrophic growth *in planta*. This assumption is supported by regulated *kpp6* expression that is induced by the b-heterodimer and Rbf1 (30, 64). Expression levels peak after plant penetration (2 dpi) and remain high during subsequent stages in the plant. In comparison, expression of *kpp2* is even slightly reduced after plant penetration (41).

In essence, the negative feedback of the UPR on the pheromone response pathway negatively regulates mating and pathogenic development prior plant infection but likely supports biotrophic growth *in planta*. We identified a UPR-regulated automatic gain control that is predicted to support efficient plant colonization, fungal biotrophy and ER homeostasis during the interaction with the host plant.

## Materials and Methods

### Strains and Growth Conditions

*Escherichia coli* TOP10 strain was used for cloning purposes and amplification of plasmid DNA. *Ustilago maydis* cells were grown at 28°C in YEPS light medium (65), complete medium (CM) (66) or yeast nitrogen base (YNB) medium (67, 68). *crg1*-driven gene expression was induced as described previously (30). Yeast-two-hybrid analysis was performed according to the Matchmaker 3 manual (Clontech). *S. cerevisiae* cells (AH109) containing Y2H plasmids were grown in SD medium -Leu/-Trp and spotted on SD solid media -Leu/-Trp or -Ade/-His/-Leu/-Trp. *U. maydis* strains used in this study are listed in table S1.

### DNA and RNA Procedures

Molecular methods followed described protocols (69). DNA isolation from *U. maydis* and transformation procedures were performed according to (70). For gene deletions, a PCR-based approach and the *SfiI* insertion cassette system was used (71, 72). Linearized plasmid DNA or PCR amplified DNA was used for homologous integration into the genome. Correct integration was verified by Southern hybridization. Total RNA was extracted from exponentially growing cells in axenic culture using Trizol reagent (Invitrogen) according to the manufacturer’s instructions. To check RNA integrity, total RNA was run on an agarose-gel and visualized by ethidium bromide staining. Residual DNA was removed from RNA samples using the TurboDNA-*free*^TM^ Kit (Ambion/Lifetechnologies). cDNA was synthesized using the RevertAid First Strand cDNA Synthesis Kit (Thermo Scientific). Primers used in this study are listed in table S2.

### Quantitative RT-PCR

qRT-PCR analyses were performed as described before (60). qRT-PCR experiments were performed with three independent biological replicates and two technical replicates each (if not stated otherwise) using the MESA Green qPCR Mastermix Plus (Eurogentec). qRT-PCR was performed using the CFX Connect Real-Time PCR Detection System and analyzed with the CFX Manager Maestro Software (BioRad).

### Plasmid Construction

For the *kpp2-GFP* fusion, 1 kb downstream of the *kpp2* stop-codon and 1 kb of the *kpp2* open reading frame (ORF) were PCR amplified from genomic DNA and ligated to the 3.7 kb *SfiI GFP-Hyg*^R^ fragment of pUMa317 (73). The resulting ligation product was integrated into the pCR2.1 TOPO vector (Invitrogen) generating plasmid pCR2.1 pUMa317-*kpp2*. For generation of the *fuz7^DD^-3xHA* fusion, the *fuz7^DD^* gene was amplified from the p123P*_crg1_:fuz7^DD^* plasmid (6) adding a C-terminal HA-tag and two *NdeI* restriction sites. The resulting 1.3 kb *NdeI fuz7^DD^-HA* fragment was ligated to the 8.1 kb *NdeI* fragment of the p123P*_crg1_:fuz7^DD^*backbone resulting in plasmid p123*fuz7^DD^-HA*. To generate the *rok1-mCherry* fusion, 1 kb downstream of the *rok1* stop-codon and 1 kb of the *rok1* ORF were PCR amplified from genomic DNA and ligated to the 3.1 kb *SfiI mCherry-G418* fragment of plasmid pUMa1723. The ligation product was integrated into the pCR2.1 TOPO vector (Invitrogen) generating plasmid pCR2.1 *rok1-mCherry*.

To generate *rok1* deletion construct, 1 kb upstream of the *rok1* start codon and 1 kb downstream of the *rok1* stop codon were PCR amplified and ligated to the 2.4 kb *SfiI Phleo*^R^ fragment of vector pUMa263.

To replace the carboxin resistance in vector P*_cib1_:cib1*^s^(18) by nourseothricin resistance, the 4.2 kb *HindIII Nat*^R^ fragment of vector pNEB-Nat^R^ and the 4.1 kb *HindIII* fragment of vector P*_cib1_:cib1*^s^ were ligated to generate the plasmid pNEB-NatR-*cib1*^s^.

To obtain the promoter fusions/gene replacement constructs for plant infection, 1 kb flanking regions up- and downstream of the genes *UMAG_12184* and *UMAG_03597*, the *kpp2* ORF and the *Hyg*^R^ of vector pUMa1442 were PCR amplified. Thereby, *SfiI* restriction sites were added to the *kpp2* gene, a *SfiI* restriction site was added to the 3’end of the upstream flanking region, a *SfiI* (N-terminal) and a *BamHI* (3’end, for *UMAG_12184*) or *KpnI* (3’end, for *UMAG_03597*) restriction site were added to the *Hyg*^R^, and a *BamHI* (5’end, for *UMAG_12184*) or *KpnI* (5’end, for *UMAG_03597*) restriction site was added for the downstream flanking region of the respective gene. The obtained fragments were ligated to obtain a *LB-kpp2-Hyg*^R^*-RB* fragment. The fragment was then integrated into the pCR2.1 TOPO vector (Invitrogen) generating plasmids pCR2.1 P*_UMAG_12184_:kpp2* (Hyg^R^) and pCR2.1 P*_UMAG_03597_:kpp2* (Hyg^R^).

For generation of the yeast-two hybrid fusion proteins, the genes of interest were PCR amplified adding *SfiI* restriction sites and ligated to modified version of the vectors pGBKT7 and pGADT7 (Clontech) containing *SfiI* restriction sites.

### Mating assay/ Filamentous growth assay

Mating assays were performed as described in (30). Cells were grown in CM-glucose medium overnight, OD_600_ was measured and adjusted to 0.2 in fresh CM-glucose medium. Cells were grown for 4 h at 28°C. OD_600_ was then adjusted to 1.0 in CM-glucose medium. Compatible FB1 and FB2 strains and derivatives were mixed 1:1 and 5 µl of the mixture was spotted on potato dextrose medium supplemented with 1% charcoal (PD-CC) (66). To test for filamentous growth of SG200, 5 µl of SG200 and derivatives were spotted on PD-CC medium and incubated for 24 to 48h at 28°C.

### Pheromone and cAMP treatment

To test for conjugation tube formation of cells in liquid culture, cultures of *U. maydis* strains were grown in CM-glucose medium at 28°C to reach an OD_600_ of 0.25. Synthetic *a2* pheromone (f.c.: 2.5 µg/ml) or DMSO as negative control was added and 2 mL of cell suspension were incubated in a 15 mL reaction tube on a rotating wheel with 6 rpm for 6 h. Cells were then microscopically analyzed using an Axio Imager.M2 equipped with an AxioCam MRm camera (ZEISS). Images were processed using ImageJ. Cells were harvested by centrifugation and flash frozen in liquid nitrogen for subsequent mRNA isolation.

For cAMP treatment, *U. maydis* strains were grown for 6 h at 28°C in CM liquid medium supplemented with 1% glucose (CM-glucose to reach an OD_600_ of 0.2 and treated with 6mM cAMP (f.c.) for 12 h at 28°C, harvested by centrifugation and flash frozen in liquid nitrogen for subsequent mRNA isolation.

### Protein procedures

For preparation of protein extracts, over-night cultures of *U. maydis* were grown in CM liquid medium supplemented with 1% glucose and adjusted to an OD_600_ of 0.25 in CM supplemented with 1% glucose (non-inducing) or arabinose (induction of P*_crg1_*-controlled expression) and further incubated for 6h at 28°C. Cells were then harvested by centrifugation and washed once with lysis buffer (50 mM Tris-HCl, pH 7.5, 150 mM NaCl, 0.1% Triton-X-100) supplemented with 2x Complete protease inhibitor (PI, ROCHE). Cell pellets were resuspended in 100-200 µl lysis buffer +2xPI, supplemented with glass beads and flash frozen in liquid nitrogen. Cell lysis was performed on a vibrax shaker (IKA) at 2200 rpm at 4°C for 30 min. The resulting suspension was cleared by centrifugation at 20000 rcf at 4°C for 10 min. The supernatant was supplemented with 4x Roti-load 1 buffer (Carl Roth), boiled for 5 min at 95°C and subjected to SDS-PAGE.

For immunoprecipitation of proteins, over-night cultures of *U. maydis* were grown in CM supplemented with 1% glucose and adjusted to an OD_600_ of 0.25 in CM supplemented with 1% glucose (non-inducing) or arabinose (induction of P*_crg1_*-controlled expression) and incubated for 6h at 28°C. Cells were then harvested by centrifugation, washed once with IP lysis buffer (25 mM Tris-HCl, pH 7.4, 150 mM NaCl, 1 mM EDTA, 1% NP-40, 5% Glycerol) supplemented with 2x Complete protease inhibitor (PI, ROCHE). The pellet was frozen in liquid nitrogen and disrupted in a cell mill (Retsch MM400, 30Hz, 2 min, 2x). The disrupted cells were then resuspended in 750 µL IP lysis buffer + 2x PI and centrifuged at 22000 rcf for 30 min at 4°C. The supernatant was added to 25 µl of agarose GFP-Trap or RFP-Trap beads (Chromotek) and incubated for 3h at 4°C on a rotating wheel. After incubation, beads were washed 3-5 times with IP lysis buffer +1x PI. 50 µL 2x Roti Load 1 (Carl-Roth) were added to the beads and boiled at 95°C for 5 mins. Samples were run on a 10% (Kpp2-GFP/Fuz7^DD^-HA) or 8% (Rok1-mCherry) SDS-PAGE and subsequently subjected to Western hybridization. All experiments were repeated at least three times.

Commercially available α-GFP (Sigma-Aldrich, 1:4000), α-HA (Sigma-Aldrich, 1:4000) or α-RFP [G6G] (Chromotek, 1:1000) antibodies were used to detect respective fusion proteins. As secondary antibodies horseradish peroxidase (HRP)-conjugated α-mouse or α-rabbit IgG (Promega, 1:4000) were used. Luminata Crescendo Western HRP substrate (Merck Millipore) was used for chemiluminescence based detection of the fusion proteins.

### Plant Infection studies

For infection studies of maize (*Zea mays*), the solopathogenic strain SG200 and derivatives or FB1 and FB2 and their respective derivatives were grown to an OD_600_ of 0.6-0.8 in YEPS light medium, adjusted to an OD_600_ of 1.0 in water. Haploid strains (FB1. FB2 and derivatives) were mixed 1:1 with a compatible mating partner. Approximately 500 µl of the resulting cell suspension were used to inoculate 8-day-old maize seedlings of the maize cultivar Early Golden Bantam. Plants were grown in a CLF Plant Climatics GroBank with a 14 h (28°C) day and 10h (22 °C) night cycle. Symptoms were scored according to disease rating criteria reported in (27). Three independent clones were used for plant infection experiments or as otherwise indicated and the average scores for each symptom are shown. Photos from infected leaves were taken und represent the most common infection symptom for the respective mutant.

### Statistical Analysis

Statistical significance was calculated using Student’s *t* test. The statistical significance of plant infection phenotypes was calculated using the Mann-Whitney test as described previously (68). Results were considered significant if the P value was <0.05.

### Accession numbers

Sequence data from this article can be found in the National Center for Biotechnology Information database under the following accession numbers:

*biz1, UMAG_02549*, XP_011388956.1; *hdp1, UMAG 12024*, XP_011391576.1; *hdp2, UMAG_04928*, XP_011391247.1; *hap2, UMAG_01597*, XP_011387600.1; *rop1, UMAG_12033*, XP_011392517.1; *mfa1, UMAG_02382*, XP_011388682.1; *pra1, UMAG_02383*, XP_011388683.1; *prf1, UMAG_02713*, XP_011389082.1; *UMAG_00306*, XP_011386216.1; *UMAG_05348*, XP_011392050.1; *UMAG_06190*, XP_011392560.1; *UMAG_10838*, XP_011389822.1; *fuz7, UMAG_01514*, XP_011387552.1; *kpp2, UMAG_03305*, XP_011389711.1; *rok1*, *UMAG_03701*, XP_011390174.1; *kpp6*, *UMAG_02331*, XP_011388645.1; *eIF2b, UMAG_04869*, XP_011391708.1

## Acknowledgements

This work was supported by funding from the Deutsche Forschungsgemeinschaft (DFG) in the framework of the IRTG 2172 PRoTECT. We thank Regine Kahmann for strains, plasmids and the generous gift of synthetic *a2* pheromone and Gerhard Braus and Ivo Feussner for generous support.

## Supporting Information

**Fig. S1: Mating is reduced in strains expressing *cib1*^s^.** Mating assay of FB1, FB2, FB1*cib1*^s^ and FB2*cib1*^s^. Mixtures of compatible haploid strains were spotted on PC-DD solid media, incubated for 36 h at 28°C and scored for filamentous growth. White fuzzy colonies indicate the formation of filaments.

**Fig. S2: UPR activity does not affect *rok1* transcript or Rok1 protein levels. (A)** qRT-PCR analysis of *rok1* gene expression in FB1*fuz7^DD^*(WT) and a derivative expressing *cib1*^s^. Cells were grown for 6 h in CM-glucose (non-inducing) or CM-arabinose (inducing) medium at 28°C. *eIF2b* was used for normalization. Expression values represent the mean of three biological replicates with two technical duplicates each. Error bars represent the SEM. **(B)** Immunoblot analysis of Rok1-mCherry protein levels. Derivatives of strains FB1*fuz7^DD^ rok1-mCherry* (WT) and FB1*fuz7^DD^ rok1-mCherry cib1*^s^ (*cib1*^s^) were grown in CM-glucose (non-inducing) or CM-arabinose (inducing) liquid medium for 6 h at 28°C. Rok1-mCherry protein was enriched from total protein extracts by immunoprecipitation with RFP-Trap beads. Total protein amount was normalized before immunoprecipitation. Rok1-mCherry was visualized using an α-RFP [G6G] specific antibody.

**Fig. S3: Rok1 physically interacts with the MAPKs Kpp2 and Kpp6.** Yeast strain AH109 was transformed with pGBKT7-Rok1 and pGADT7-Kpp2 or pGBKT7-Rok1 and pGADT7-Kpp6 and spotted on SD-Leu/-Trp and SD-Ade/-His/-Leu/-Trp solid media. AH109 transformed with pGBKT7-p53 and pGADT7-T or pGBKT7-lam1 and pGADT7-T (Clontech) served as positive and negative control, respectively. Yeast strain AH109 was transformed with pGBKT7-Rok1 and pGADT7 and spotted on SD-Leu/-Trp and SD-Ade/-His/-Leu/-Trp solid media to test for auto-activation of Rok1.

**Table S1: Strains used in this study**

**Table S2: Primers used in this study**

