## Supplementary figures and images for "The unfolded protein response regulates pathogenic development of *Ustilago maydis* by Rok1-dependent inhibition of mating-type signaling"

### Fig. S1

**S1**

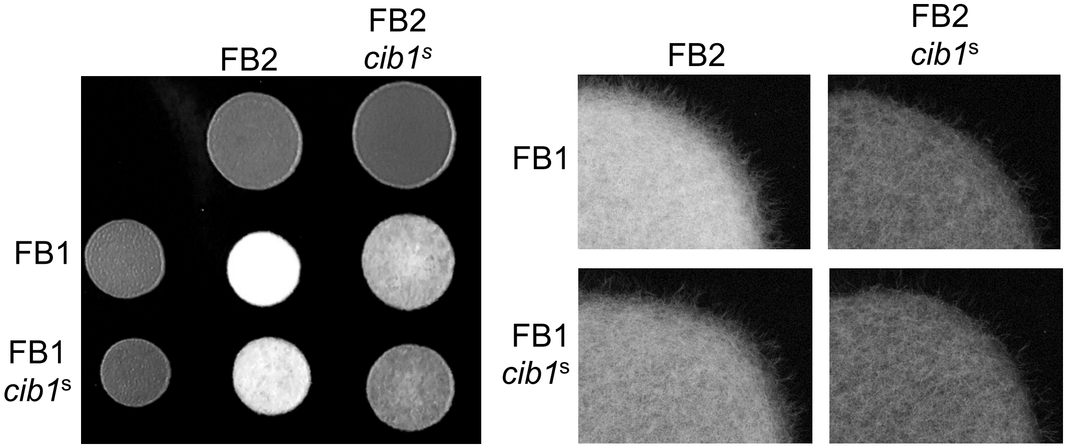

### Fig. S2

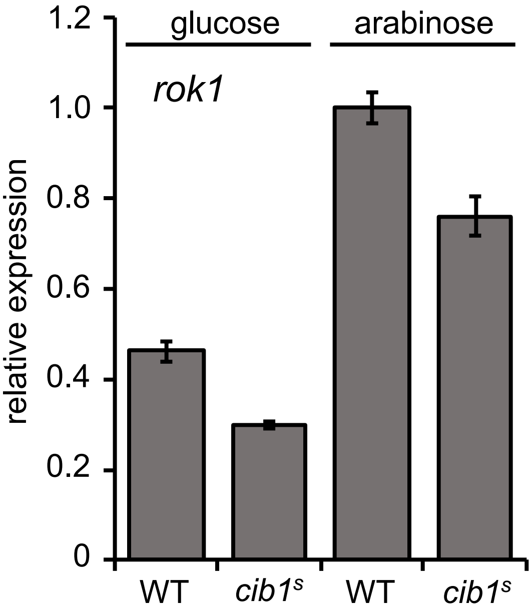

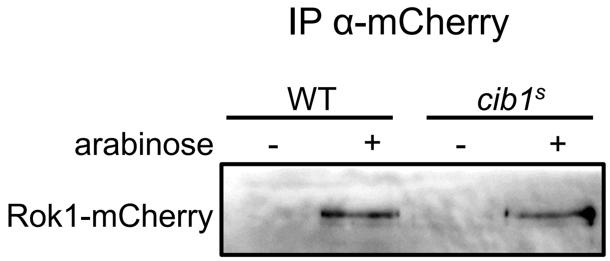

### Fig. S3

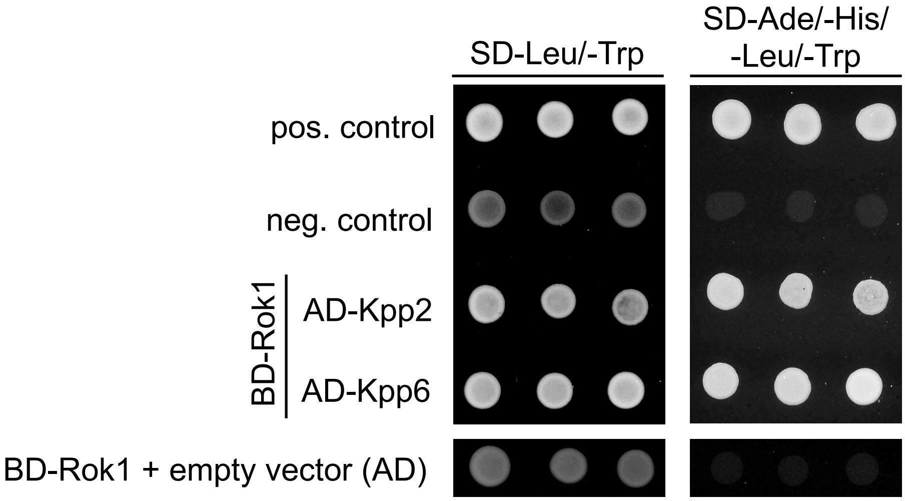
